# Landscape conservation and orchard management influence carob tree yield through changes in pollinator communities

**DOI:** 10.1101/2024.07.05.602169

**Authors:** Carmelo Gómez-Martínez, Miguel A. González-Estévez, Indradatta deCastro-Arrazola, Peter Unglaub, Amparo Lázaro

## Abstract

Worldwide pollinator declines are a major problem for agricultural production. However, understanding how landscape characteristics and local management influence crop production through its pollinators is still a challenge. By sampling 20 orchards in Mallorca island (Spain), we evaluated how the landscape (habitat loss) and orchard local management (farming system: conventional vs. ecological; male-to-female ratio) influenced pollinator communities and production in carob trees (*Ceratonia siliqua*), a crop of high economic importance in Mediterranean areas. We found that orchards surrounded by larger natural areas received more visits by wild bees and butterflies and fewer by honeybees. Ecological farming tended to increase overall pollinator abundance in the orchards. High male-to-female ratio enhanced overall pollinator abundance and shaped pollinator composition, by increasing hoverfly abundance and decreasing wasps and flies. Male-to-female ratio showed hump-shaped relationships with fruit and seed production per female tree (peak at 0.7 males/female), whereas total orchard production maximized with 20-30% of males. Seed weight (farmer’s highest economic value) increased in conserved landscapes where wild pollinators prevailed, and with overall pollinator abundance; however, it decreased with male-to-female ratio, likely due to seed number-size trade-offs. Management strategies to enhance carob production may optimize sex ratios and favor wild pollinators by reducing pesticide use and preserving natural landscapes.

## Introduction

The ongoing decline of wild insects [1–3] supposes a threat to the essential ecosystem service of pollination [4]. Land-use changes are considered one of the main factors driving such pollinator decline, as they lead to the homogenization of landscapes [5] and communities [6], affecting habitat and resource availability for pollinators [7], and disrupting plant-pollinator interactions [8]. This pollinator loss is especially worrisome for the maintenance of agricultural production, because the 35% of cultivated plants depend on insect pollination [4]. Although managed honeybees are commonly used to pollinate crops, wild insects are often more effective pollinators [9–11], and pollination by wild insects is known to increase crop production quantity [4, 10, 12] and quality [13] and therefore, farmers’ profits [14].

Several studies have shown that habitat loss has a negative effect on pollinator visits and production in crops (e.g., [12, 15–18]). A larger amount of natural habitat in the landscapes allows a richer insect interchange between natural habitats and crops [19], while landscapes dominated by crops may have abundant but little diverse pollinator communities [20]. Besides, local crop management may have additional effects on its pollinators and production. A higher pollinator diversity and pollination success has been found in ecological crops [21–24] and in orchards with a living ground cover [15, 25], which is often part of ecological or traditional farming systems. Besides, the negative effects of non-ecological management on pollinator abundance and richness may be stronger in simplified than in complex landscapes [26, 27].

Lastly, the effect of pollinator loss on crops might depend on crop breeding system and its dependence on pollinators [28–30]. Particularly, dioecious crops, which rely on pollinator visitation to both sexes for reproduction, might be particularly vulnerable to overall pollinator decrease [31], as well as to the loss of the most efficient ones [32]. For this type of crops, optimizing the sex ratio during orchard design could be critical [33]. An equilibrated ratio might be important to optimize crop production, not only because more females imply more individuals producing seeds, but also to control for density-dependent processes of facilitation-competition between sexes [34]. Indeed, a too high proportion of males might be counterproductive for pollinator visits to females in cases where males are more attractive to pollinators [35–37].

The carob tree (*Ceratonia siliqua* L.) is a perennial crop extended in the Mediterranean climate region [38, 39]. It is a polygamotrioecious species, although it mainly behaves as dioecious [40–42], and depends on insect pollinators for reproduction [43]. Spain is the main producer of carob for commercial purposes with around 60-80 thousand tons per year [44]. Although it was traditionally used as cattle food, nowadays carob cultivation is uprising due to the use of seeds and pulp in food and pharmaceutical industries [45–48]. As a result of this industrial interest, most of the studies on carob trees have focused on chemical and pharmaceutical properties (e.g., [48–52]). To our knowledge, there are no studies to date that evaluate how the production of this important crop is influenced by landscape and local effects on its pollinators. Understanding how the landscape context and local management of orchards affect carob tree yields is important to palliate the potential negative effects of landscape disturbances on wild pollinator communities, and to enhance crop production in a sustainable manner. With this aim, we collected data on pollinator visitation to carob tree flowers and fruit and seed production in 20 carob tree orchards along a gradient of habitat loss in Mallorca Island (Spain). Particularly, we analyzed how the percentage of surrounding natural habitat (Fig. 1A), the farming system (conventional vs. ecological; Fig. 1B) and the ratio of male-to-female trees in the orchards (Fig. 1C) influenced: 1) the composition and abundance of pollinators visiting carob orchards; 2) the production of carob fruits and seeds, and 3) the quality of carob seeds, estimated as seed weight. We expected that orchards within conserved landscapes and subjected to ecological farming held a higher pollinator abundance and community compositions that enhanced carob production; and that the negative effect of habitat loss on crop production was less pronounced in ecological farms. Lastly, we expected an increase in pollinator visitation rates with the increase in male trees in the orchards, whereas a hump-shaped relationship between the proportion of male trees and crop production related to density-dependent competitive processes between sexes.

**Fig. 1.**
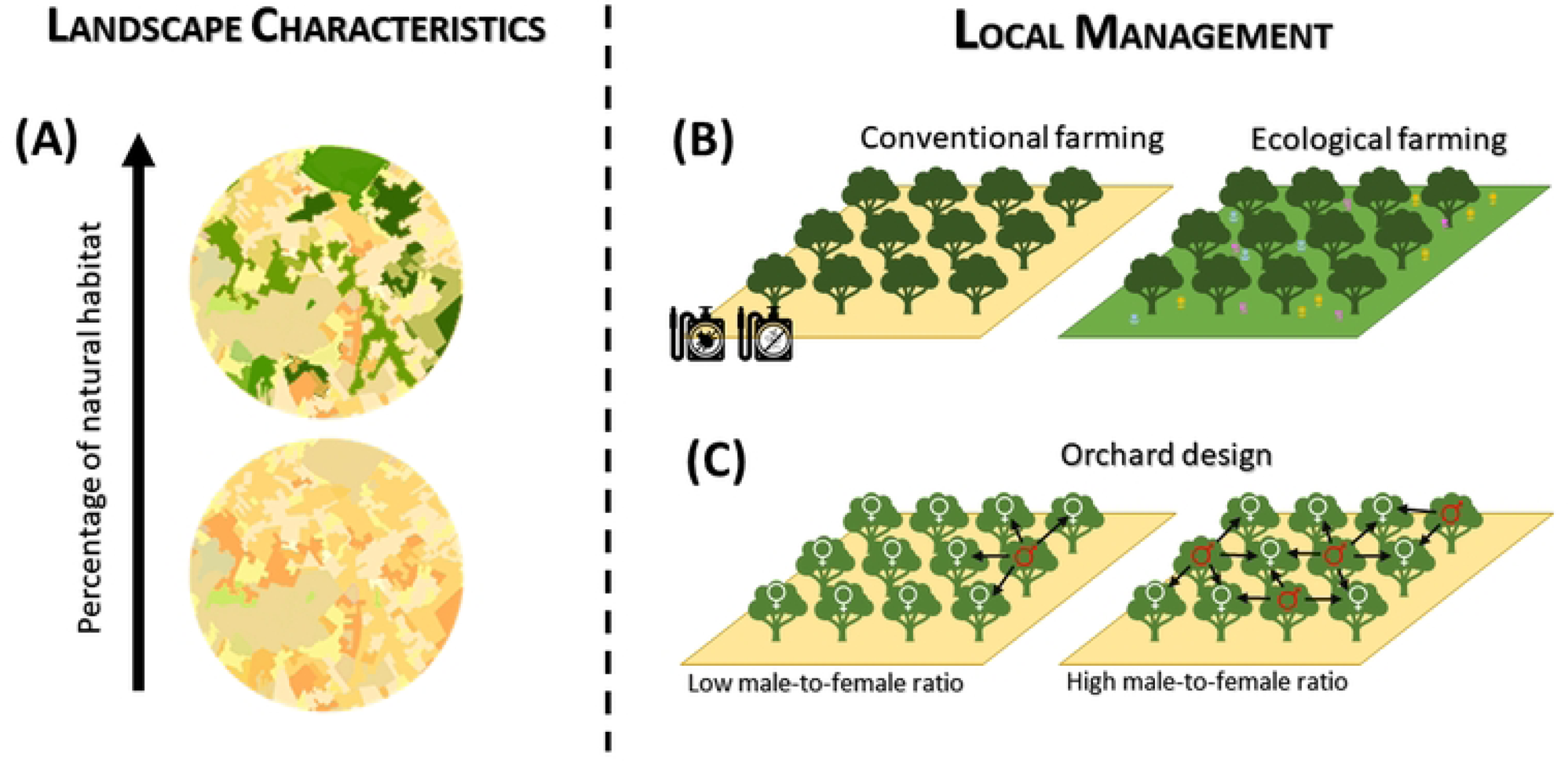
Conceptual diagram with the variables potentially influencing carob tree production. (A) The percentage of natural habitats in the landscape surrounding the orchards; (B) The farming system (Conventional vs Ecological Farming); (C) The proportion of carob male trees per female tree.

## Material and Methods

### Study crop

The carob tree (*Ceratonia siliqua* L.) is a crop belonging to the family Fabaceae, widely cultivated in the Mediterranean region, and introduced in other areas with Mediterranean climate, such as California, Australia, or South Africa [38, 39]. The carob pods, seeds, and gum have many nutritional, biochemical, and clinical applications, as for instance for livestock feeding, in food and beverages, bioenergy production, in pharmaceutical industry in pomades, anti-celiac ingredients, pills, and dental paste [45–48]. Spain is the first producer worldwide (60-80 thousand tons per year) [44], and the Balearic Islands concentrate around a third of the Spanish production. Carob market prices are cyclic, with long gaps and short high peaks that happened on 1984, 1994, 2006, 2011 [39], and recently in 2022, when the prices of the Spanish and Portuguese production in the international market raised from 0.8€/kg to more than 2€/kg for whole pods and from 7€/kg to 27€/kg for seeds [53].

Carob tree is a polygamotrioecious species, but it mostly behaves as dioecious, with around 50% of wild trees being males, 49% females, and only 1% hermaphrodites [40–42]. The flowers are inconspicuous, with no developed corolla, and grow in racemes in old branches. The fruits are dark-brown pods with hard seeds and a sweet pulp and take almost a year to mature [54]. Carob trees are highly dependent on animal pollination for reproduction [55]. Male trees are more fragrant and tend to have a larger flower display, which could serve as attractant for pollinators [38, 43]. However, as male trees do not produce fruits, its abundance in crops is usually much lower than the abundance of female trees, or even null, which might drive to a spatial isolation of female trees, hindering pollination [56].

### Study orchards

We selected 20 carob tree orchards (study orchards, hereafter) across Mallorca Island (Fig. S1 in Supporting Information), the main island of the Balearic Archipelago (Spain). The study orchards were selected along a gradient of natural habitat loss (see below), and separated from each other by a minimum distance of 1.5 km from the center the orchards (mean ± SE distance among study orchards: 32.07 ± 1.42 km). All the study orchards flowered during the same period (from beginning of October to mid-end November), however, it cannot be totally discarded that they include different cultivars, as this information was unknown by the farmers.

### Landscape characterization

We used the last update of the SIOSE AR database (Spanish acronym for Soil Occupation Information System of Spain - High Resolution) [57] to describe the characteristics of the landscape surrounding the study orchards within a buffer zone of 1-km radius from their center. We had a total of 43 land-cover classes in the buffer zones, 7 of which corresponded to natural and semi-natural habitats including pastures, mix of shrub and pasture, mix of wood and pasture, woodland, perennial and coniferous forest, and shrubland. The remaining 36 land-cover classes corresponded to crops (mainly non-citric fruit trees and herbs; 13 classes), artificial (17 classes) and water bodies (6 classes). With these data we estimated the percentage of natural areas from the total area within the 1-km buffers (% natural habitat, hereafter; Table S1 in Supporting Information).

### Local management of orchards

To study potential effects of the local management on the pollinator community and production of carob tree orchards, we considered the farming system (conventional vs. ecological), and the male-to-female ratio (number of male trees per female tree). We classified the study orchards as conventional or ecological according to the European regulations, which in the Balearic Islands are managed and applied by the CBPAE (acronym in Catalan for the Council for Ecological Agricultural Production of Balearic Islands). Seven of the study orchards were certified as ecological farms, while the remaining 13 were not (Table S1). According to the CBPAE, ecological farming does not make use of synthetic agrochemicals, maintains non-cultivated areas and natural habitats to promote biodiversity, must be sustainable, and do not use genetically modified organisms (GMO). Regarding the male-to-female ratio, we registered the sex of every carob tree in each study orchard and divided the number of males by the number of females to get a ratio of males to female (male-to-female ratio, hereafter; Table S1). The male-to-female ratio in carob orchards of Mallorca is generally low, as farmers obtain female carob trees by grafting female branches in male rootstocks to increase the number of producer trees. Hermaphrodite trees were absent in 11 of the 20 study orchards and scarce in the rest (from 1 to 8%), except for one study orchard where most of the individuals (95.7%) were hermaphrodite (Son Cotoner; Fig.S1). For this reason, and to simplify the male-to-female ratio estimate in the study orchards, we counted the hermaphrodite trees both as males and females.

### Pollinator visitation

We recorded visits of pollinators to carob tree flowers during two flowering seasons (2019 and 2020), from the beginning of October to the beginning of November, covering the flowering period of this crop in Mallorca. Each study orchard was sampled five days during the flowering period (approximately once per week). We observed pollinator visits to carob flowers while we walked slowly along tree lines for one hour each sampling day at each study orchard. Pollinator censuses were carried out between 10.00h and 16.00h, in sunny days without wind, to ensure pollinator activity. We considered a visit of a pollinator to a flower when a pollinator was observed contacting the reproductive parts of the flower. We noted if the pollinator contacted a male, female, or hermaphroditic flower. As flowers are very close to each other in a raceme, we counted as one interaction a visit of a pollinator to a raceme, although they might contact several flowers. As flower racemes of carob trees are caulogenous (emerging only from the old branches) [58], collecting visiting insects between branches and leaves with a hand-net becomes difficult. Therefore, during censuses, we categorized each visitor into eight functional groups: butterflies, hoverflies, bee flies, other flies (mainly muscoids and small acalyptrate flies), ants, wasps, wild bees, and honeybees (*Apis mellifera*). Nevertheless, we captured the insects when possible for their identification in the lab if their identification to the species level in the field was unfeasible (pausing the watch during manipulation to standardize the time spent per study orchard). We collected 70 specimens in 2019 and 44 in 2020 (Table S2 in Supporting Information for species list). With these data we calculated (1) the pollinator abundance by functional group, as the total number of individuals of each functional group recorded in each study orchard each year, and also (2) the overall pollinator abundance, as the total number of individuals recorded in each study orchard each year.

### Carob tree production

Both study years, at the beginning of the flowering period (beginning of October in 2019 and 2020), we randomly selected 20 female trees in each study orchard and haphazardly marked one of its branches to count the number of inflorescences and inflorescence buds. We then counted the number of flowers and flower buds in five random inflorescences of each marked branch and estimated the total number of flowers in the marked branch as the average number of flowers per inflorescence multiplied by the number of inflorescences in the branch. After fructification (August in 2020 and 2021) and before harvest, we counted the number of aborted and developed fruits in each marked branch, and estimated fruit set in them as the number of developed fruits divided by the total number of flowers. We also collected five fruits per marked branch to study seed production and weight in the lab, by counting the number of developed and aborted seeds separately for each of the five fruits per tree, and weighting all the developed seeds per fruit together.

With all these data, we then calculated (1) *Fruit production per female tree*, as the fruits produced by 1000 flowers in a tree (fruit set x 1000); (2) *Seed production per female tree*, as the seeds produced by 1000 flowers in a tree (seeds/fruit x fruit production per tree); and (3) *Seed weight*, as the weight (in grams) of one seed, calculated for each fruit as the total mass of developed seeds divided by the number of developed seeds in that fruit. Fruit and seed production per tree were estimated per 1000 flowers because fruit set is very low in carob trees, but the number of fruits per tree may be relatively high due to the large number of flowers produced [54].

To shed light on the optimum male-to-female ratio in carob tree orchards, we additionally estimated the percentage of males in the orchards (males/100 trees) that maximized total fruit and seed production, and the total weight of seed production at the orchard level (hereafter orchard-level fruit production, orchard-level seed production, and orchard-level total seed mass, respectively). We estimated orchard-level fruit and seed production by multiplying the production of a female tree by the percentage of females in the orchard. Orchard-level total seed mass was estimated as the orchard-level seed production multiplied by the estimated seed weight at different male-to-female ratios (see below for results of GLMMs). Note that fruit and seed production per tree were estimated per 1000 flowers.

### Statistical analysis

We performed all the statistical analyses in R v.4.2.2 [59]. To study how pollinator community composition changed with the percentage of natural habitat surrounding the study orchards, the male-to-female ratio and the farming system (conventional vs ecological), we run a Canonical Correspondence Analysis (CCA; R-package vegan v.2.6.4) [60]. We also included the year as predictive variable, as pollinator communities might change between study years. The response variables were the abundance (number of visits) per study orchard and year of each pollinator group: honeybees, wild bees, wasps, ants, hoverflies, other flies, and butterflies. As the number of beetles and bee flies was extremely low (see Results section), we removed the beetles and the bee flies from the analyses. To assess the relationship between the response variables and the predictors, we first evaluated the significance of the whole ordination by performing Monte Carlo permutations [60, 61]. The significance of the model was determined by comparing the observed statistics against a null distribution generated from 9999 random permutations of the community data, while keeping the predictors fixed. To determine the significance of each predictive variable, we performed conditional permutation tests. In these tests, the significance of each variable was evaluated by randomizing the variable to test while keeping the rest of predictors fixed. The observed statistics for each variable were compared against a null distribution generated from 9999 random permutations [60, 61].

To evaluate how overall pollinator abundance was related to the surrounding landscape and the local management of the study orchards (male-to-female ratio and farming system), we fitted a generalized linear mixed model (GLMM; R-package lme4 v.1.1.32) [62], with the study orchard as a random factor to control for pseudoreplication. In this model, we included the percentage of natural habitat, the male-to-female ratio, the faming system (conventional vs. ecological) and the sampling year as predictor variables in the full model, after checking for the absence of collinearity (VIF values < 3) [63]. As previous studies showed a potential synergic effect between landscape disturbance and local management [26, 27], we also included the interaction between the percentage of natural habitat and the farming system in the full model. To evaluate how carob trees’ production was related to the landscape surrounding the study orchards and the local management, we also fitted separated GLMMs with fruit production per female tree, seed production per female tree, and seed weight as response variables. The predictor variables were the same as for the GLMM models of pollinator visits, but we also added total pollinator abundance and the first axis of the CCA as predictor variables to assess whether the abundance and composition of pollinators in the study orchards directly influenced carob production. We did not include the second axis of the CCA as it was not significant (see Results section). As we expected a hump-shaped relationship between carob production and the male-to-female ratio due to density-dependent competence/facilitation processes [34, 64], we included the male-to-female ratio both as linear and quadratic terms, first by standardizing the linear term to □ = 0 and σ = 1, and then calculating its quadratic form [65] to avoid collinearity [63]. Both study orchard and tree nested within study orchard were included as random factors in these models. Due to overdispersion [63], we used a negative binomial distribution (link log) for the models of pollinator abundance, and fruit and seed production per female tree. A gamma distribution (link log) was used for the model of seed weight as it did not fulfilled the assumption of normality even after log-transformation (R-package nortest v.1.0.4) [66]. We conducted multi-model inference based on AICc (R-package MuMIn v.1.47.5) [67]. For each full model, we constructed sub-models containing different combinations of predictor variables, limiting the maximum number of predictor variables to four in each model, to avoid over-parametrization due to sample size. We then calculated unweighted averages using the sub-models with ΔAICc ≤ 2 (function model.avg from R-package MuMin v.1.47.5) [67]. Residual inspection indicated adequate model fits. Plots were drawn using the R-package ggplot2 [68], except the CCA that was drawn using the plot.cca function in R-package vegan v.2.6.4. [60].

## Results

We registered a total of 8790 pollinator visits to carob trees (4014 in 2019 and 4776 in 2020). More than half of the visits (ca. 57%; 5017 visits) were conducted by *Apis mellifera* (1857 visits in 2019 and 3160 in 2020), followed by wasps (1527 visits), muscoids and small acalyptrate flies (other flies, 917), hoverflies (561), wild bees (382), ants (241), butterflies (119), beetles (15) and bee flies (11). Table S3 in Supporting Information shows detailed information on the number of visits per pollinator group, study orchard and year. We identified 45 species of wild insects visiting carob tree flowers (Table S2), belonging to ants (6), wild bees (7), wasps (3), muscoids and small acalyptrate flies (15), hoverflies (8), bee flies (1), butterflies (4) and beetles (1). Fruit production per female tree varied between 8.8 and 45.2 fruits in one thousand flowers, depending on the study orchard (mean ± SE: 26.7 ± 2.5 fruits in one thousand flowers). Seed production per female tree varied between 88.6 and 558.3 seeds in one thousand flowers, depending on the study orchard (286.6 ± 30.35 seeds in one thousand flowers). Seed weight varied between 0.09 and 0.33 grams depending on the study orchard (0.19 ± 0.0009 grams per seed).

### Pollinator community composition

The CCA indicated significant relationships between the abundance of the pollinator groups, the landscape characteristics and local management of the orchards (*F* = 3.30, p = 0.0003; Fig. 2A). The cumulative percentage of variance explained by the first axis was 66.8%, while the two first axes explained 89.0% of the total variance. The percentage of natural habitat, the farming system and the year varied along the first ordination axis, which was significant (CCA1, *F* = 8.810, p = 0.0001). The male-to-female ratio varied along the second axis; however, this second axis was not significant (CCA2, *F* = 2.922, p = 0.1664). The predictor variables that significantly influenced the pollinator community composition were the percentage of natural habitat (*F* = 2.92, p = 0.020), the male-to-female ratio (*F* = 3.27, p = 0.018) and the year (*F =* 5.42, p = 0.001). Higher percentage of natural habitat was mainly related to an increase in the abundance of wild bees, but also of ants and butterflies, whereas more disturbed landscapes (lower percentage of natural habitats) were related to an increase in the abundance of managed honeybees. Higher male-to-female ratio was positively related to increased hoverfly abundance, while other groups as ants, wasps and other flies were related to low male-to-female ratios. Lastly, the abundance of wild bees and butterflies were higher in 2019 than in 2020, while the other pollinator groups did not significantly change their abundances between years.

**Fig. 2.**
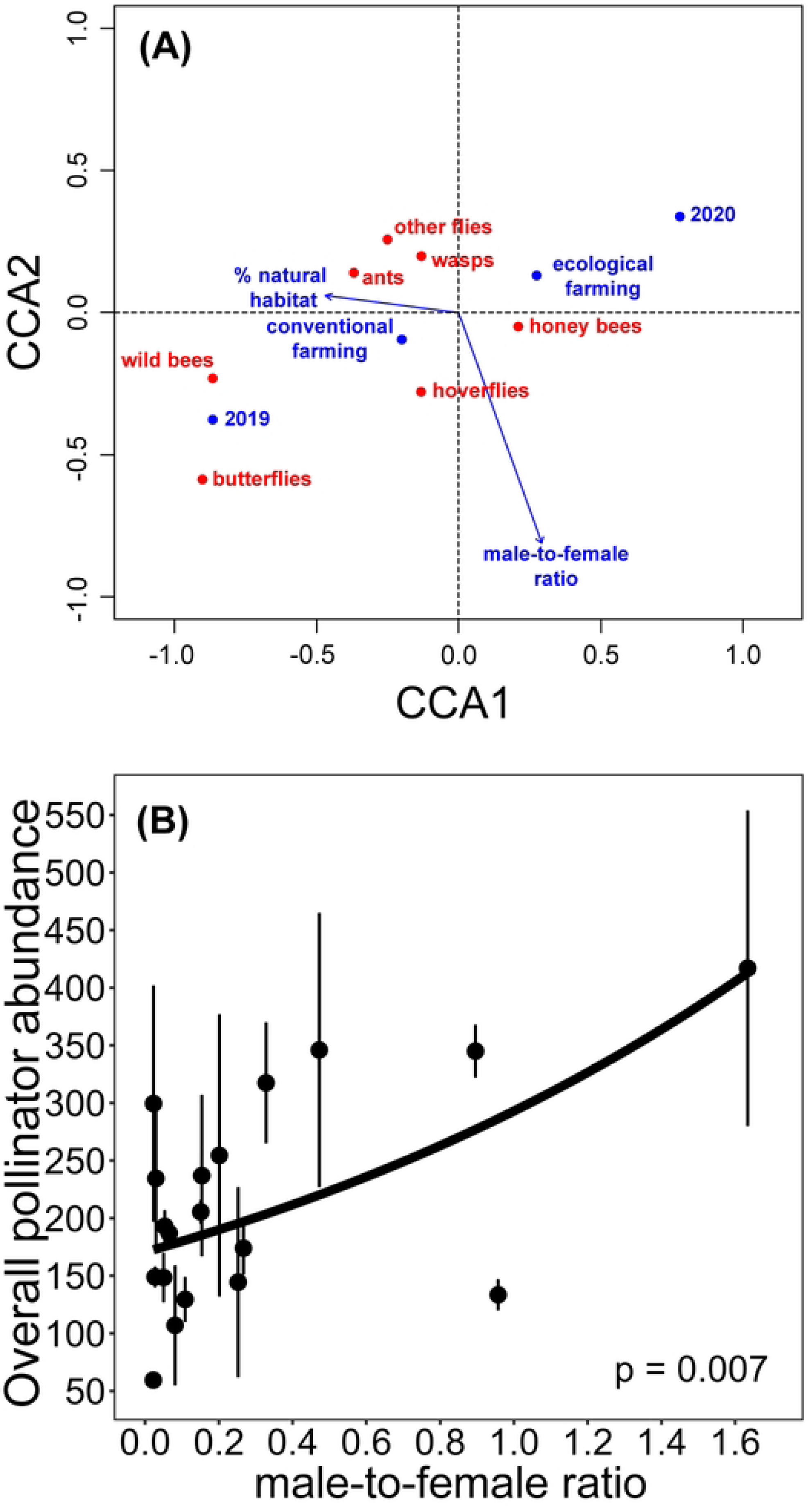
Effects of landscape and local management on overall pollinator abundance and community composition in carob orchards. (A) Canonical Correspondence Analysis (CCA) showing the relationships between the percentage of natural habitat, male-to-female ratio, farming system and year (blue arrows and dots) and the pollinator groups (red dots). Short distances between the pollinator groups and the predictor variables in the ordination indicate high association between them. Percentage of natural habitat (*F* = 2.915, p = 0.020); male-to-female ratio (*F* = 3.273, p = 0.018); year (*F =* 5.419, p = 0.0006). (B) Relationship between the male-to-female ratio and overall pollinator abundance. Lines represent the estimates of the best model, the dots represent the average of total visits per year and study orchard, and the vertical lines the standard errors.

### Overall pollinator abundance

Higher male-to-female ratios were positively related to the overall abundance of pollinators in the orchards (Table 1A, Fig. 2B). Additionally, ecological farming tended to enhance overall pollinator abundance, although the effect was only marginally significant (Table 1A).

**Table 1.** Model-averaged coefficients, significance, and relative importance (RI) of the predictor variables for the GLMMs to evaluate effects on: (A) overall pollinator abundance; (B) fruit production per female tree, (B) seed production per female tree; and (C) seed weight. Farming system refers to ecological vs. conventional practices; CCA1 refers to the first axis of the CCA and depicts the relationship between the % of natural habitats in the landscape and the composition of the pollinator community (increase abundance of wild bees with the increase in natural habitats); Male-to-female ratio and Male-to-female ratio^2^ indicate linear and quadratic terms, respectively. Significant *p*-values are marked in bold. RI was estimated by summing the Akaike weights over of all the models where the variable appears. Explanatory variables are marked in bold if RI > 0.6, which highlights the importance of these variables in the models [107].

| **Model** | **Predictor** | **Estimate** | **Std. error** | **z value** | **p** | **RI** |
| --- | --- | --- | --- | --- | --- | --- |
| (A) Overall pollinator abundance | (intercept) | 5.0921 | 0.158 | 31.334 | **< 0.0001** |  |
|  | % Natural habitat | -0.0046 | 0.005 | 0.953 | 0.341 | 0.1 |
|  | Male-to-female ratio | 0.5298 | 0.209 | 2.445 | **0.015** | **1.0** |
|  | Farming system | 0.2943 | 0.171 | 1.659 | 0.097 | 0.5 |
|  | Year | 0.1274 | 0.131 | 0.935 | 0.350 | 0.3 |
| (B) Fruit production per female tree | (Intercept) | 3.4563 | 0.170 | 20.307 | **< 0.0001** |  |
|  | CCA1 | -0.0083 | 0.056 | 0.148 | 0.882 | 0.20 |
|  | Overall pollinator abundance | -0.0006 | 0.001 | 1.064 | 0.287 | 0.21 |
|  | Male-to-female ratio<sup>2</sup> | -1.1635 | 0.525 | 2.214 | **0.027** | **1.0** |
|  | Male-to-female ratio | 0.8639 | 0.429 | 2.007 | **0.045** | **0.84** |
|  | Farming system | 0.0898 | 0.190 | 0.470 | 0.638 | 0.14 |
|  | Year | -0.1185 | 0.084 | 1.405 | 0.160 | 0.50 |
| (C) Seed production per female tree | (Intercept) | 5.8779 | 0.225 | 26.049 | **< 0.0001** |  |
|  | CCA1 | 0.0653 | 0.076 | 0.852 | 0.394 | 0.18 |
|  | Overall pollinator abundance | -0.0007 | 0.001 | 1.042 | 0.298 | 0.20 |
|  | % Natural habitat | -0.0019 | 0.006 | 0.299 | 0.765 | 0.08 |
|  | Male-to-female ratio<sup>2</sup> | -1.1711 | 0.552 | 2.119 | **0.034** | **0.92** |
|  | Male-to-female ratio | 0.8837 | 0.469 | 1.881 | 0.060 | **0.80** |
|  | Farming system | 0.1447 | 0.201 | 0.717 | 0.473 | 0.10 |
|  | Year | -0.1863 | 0.110 | 1.691 | 0.091 | **0.89** |
| (D) Seed weight | (Intercept) | -1.6990 | 0.021 | 80.691 | **< 0.0001** |  |
|  | CCA1 | -0.0219 | 0.003 | 7.028 | **< 0.0001** | **1.00** |
|  | Overall pollinator abundance | 0.0002 | 0.000 | 4.677 | **< 0.0001** | **1.00** |
|  | % Natural habitat | -0.0006 | 0.001 | 0.906 | 0.365 | 0.20 |
|  | Male-to-female ratio<sup>2</sup> | 0.0047 | 0.029 | 0.163 | 0.870 | 0.15 |
|  | Male-to-female ratio | -0.1176 | 0.040 | 2.938 | **0.003** | **1.00** |
|  | Farming system | 0.0103 | 0.027 | 0.380 | 0.704 | 0.14 |
|  | Year | -0.0022 | 0.008 | 0.273 | 0.785 | 0.14 |

### Carob tree production

The GLMMs for fruit and seed production per female tree showed similar results. Both models indicated that fruit and seed production per female tree increased with male-to-female ratio until a maximum at values of 0.6-0.7 males/female, after which higher ratio values decreased the production per female tree (Table 1B and C, Fig. 3A and B). With these data, we estimated that orchard-level fruit and seed production was maximized with 20-30% of male trees in the orchards (Fig. 3C and D). Regarding seed weight, we found a negative significant relationship between seed weight and the first axis of the CCA (CCA1, related to the percentage of natural habitat in the landscape and the abundance of wild bees vs. honeybees, Table 1D, Fig. 4A). Thus, an increase in the abundance of wild bees in more conserved landscapes was positively related to seed weight (Table 1D, Fig. 4A). The model also showed a positive significant relationship between seed weight and overall pollinator abundance (Table 1D, Fig. 4B), and a negative relationship between seed weight and the male-to-female ratio (Table 1D, Fig. 4C). Same as for fruit and seed production, we calculated that orchard-level total seed mass was maximized with 20-30% of male trees in the orchards (Fig. 4D).

**Fig. 3.**
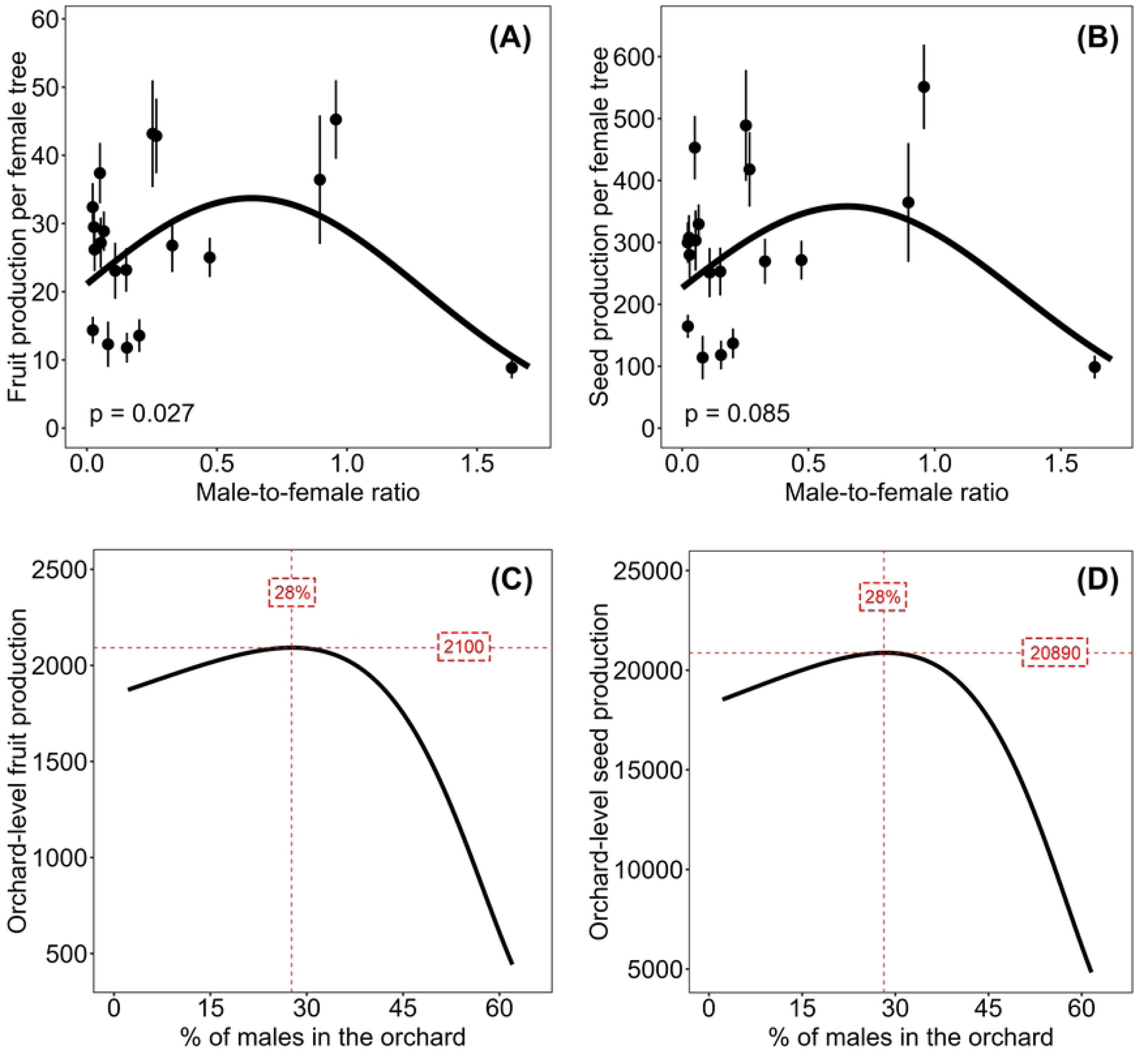
Effects of local management of orchards on the production of fruits and seeds. Panels (A) and (B) show the relationships between the male-to-female ratio and fruit and seed production per female tree, respectively. In these panels, lines represent the estimates of the averaged best models, the dots represent average fruit or seed production per year and study orchard, and the vertical lines the standard errors. Panels (C) and (D) show the relationship between the percentage of male trees (the number of male trees per 100 trees) in an orchard and the estimated orchard-level fruit and seed production, respectively (i.e., for all the females in an orchard of 100 trees). Orchard-level fruit production (or seed production) was calculated as the number of fruits (or seeds) produced by 1000 flowers in each tree multiplied by the percentage of females (the number of female trees in an orchard of 100 trees).

**Fig. 4.**
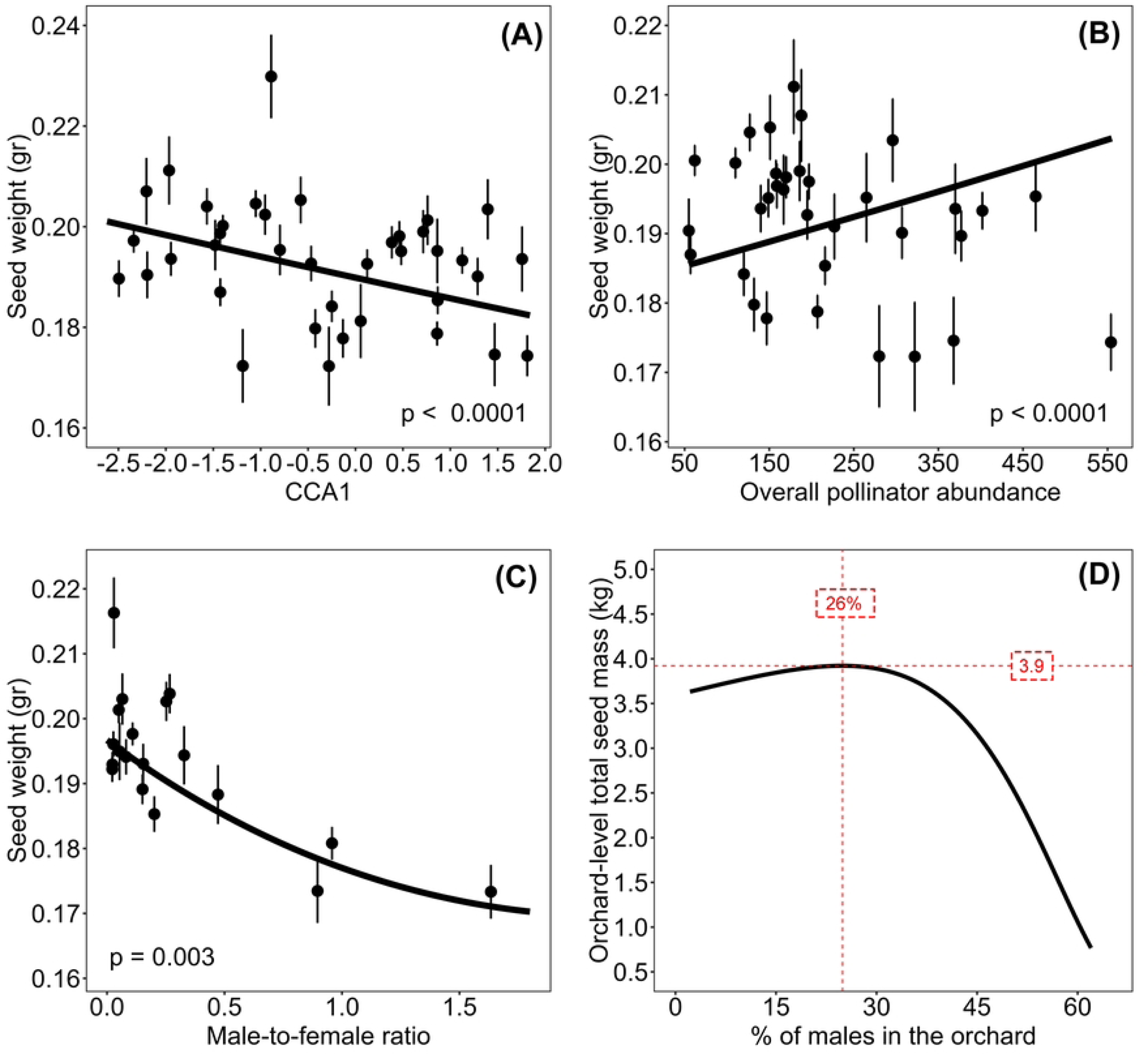
Pollinator community and the local management on seed weight. Relationships between seed weight and: (A) the first axis of CCA (CCA1); (B) Overall pollinator abundance; and (C) male-to-female ratio. Lines in panels (A-C) represent the estimates of the averaged best models, the dots represent the average seed weight per year and study orchard, and the vertical lines the standard errors. Positive values of first axis of CCA (CCA1) represent pollinator community composition related to habitat loss (mainly honeybees), while negative values represent pollinator community composition related to conserved landscape (mainly wild bees and butterflies). (D) shows the relationship between orchard-level total seed mass and the percentage of male trees (the number of male trees per 100 trees) in an orchard. Orchard-level total seed mass was calculated as the total weight of the seeds produced by 1000 flowers in each tree multiplied by the percentage of females (the number of female trees in an orchard of 100 trees).

## Discussion

Here we showed that both habitat loss and the local management (male-to-female ratio and farming system) influenced the pollinator communities of carob tree orchards. Such changes in pollinator communities directly translated into differences in carob yields, so that carob production could be maximized in orchards located within natural landscapes, subjected to ecological farming, and designed with a 20-30% of male trees.

### Pollinator communities in carob tree orchards

Honeybees were the most frequent visitor in both years, however, we found around 70% more honeybees in 2020 than in 2019. Large differences between years in the abundance of honeybees were also reported by [43], where honeybees went from being the most abundant visitor one year to be almost absent two years later. This high variation in honeybee abundance might result of their management, as the installation and removal of hives that hold several tens of thousands of individuals [69] might deeply modify their abundance in crop fields. Wasps and flies (mainly muscoids and small acalyptrate) followed honeybees as the second and third most abundant groups both years, agreeing with [58] and [70] which found wasps and Diptera as the main visitor groups together with honeybees in orchards of Spain and Israel, respectively. Only [70] identified most of the visitors to the genus or species level, naming three bee species, two wasps, one hoverfly and two flies. Although we only captured a small fraction of all the pollinators we recorded (just 115 out of the 3773 non-honeybee visitors), we were able to identify, by observation or in the laboratory, at least 45 species of wild insects (including at least five of the species identified by [70]) which makes this study the most extent census of pollinators of carob trees until date.

Most wild pollinators were linked to conserved landscapes, while honeybees were favored in more disturbed areas. These conserved landscapes offer diverse nesting sites and food resources for wild pollinators [71–73]. Pollinators with small foraging ranges like solitary bees and ants require food sources near their nests [74, 75], and even long-distance fliers typically forage close to home [76, 77]. Hence, natural habitats near the study orchards likely benefited the presence of wild pollinators [78, 79]. On the other hand, honeybees showed affinity for disturbed habitats, which might be due to the location of beehives in crop fields [80]. The male-to-female ratio also influenced the composition of pollinator communities. We found a higher number of hoverflies linked to orchards with a higher male-to-female ratio, while the abundance of wasps increased with a lower male-to-female ratio. This was expected because hoverflies feed on both pollen and nectar [81, 82], while wasps as *Vespula germanica* (the most common wasp found in our orchards) visit flowers for nectar [83].

Apart from the effect of changes in community composition, we found an overall higher abundance of pollinators in orchards with higher male-to-female ratios. This agrees with previous studies that showed male flowers to be more attractive than female flowers [43, 70, 84]. Likely this is because male trees produce a much larger number of inflorescences than female trees, have a more intense scent and offer two rewards: nectar and pollen [70, 85]. Ecological farming also tended to promote a higher pollinator abundance compared to conventional farming, but best models indicated no interaction between habitat loss and farming system. Different studies found positive effects of ecological farming on pollinator abundance and diversity [21, 23, 25], and larger positive effects of ecological farming in more disturbed landscapes [26, 27]. In our study, the difficulty of finding ecological orchards that met our landscape needs generated an unbalanced number of conventional vs ecological orchards that might have affected the detection of significant relationships. Future work with a larger number of ecological fields along landscape gradients may help test further these relationships.

### Carob tree production

We recorded a relative low fruit set compared with the high number of flowers that carob trees produce, which is not rare as the surplus of flowers may be just a mechanism to attract pollinators [86, 87]. In addition, fruit production per female tree was lower in 2020 than 2019, which could be due to biennial bearing [38, 88, 89], but also to the lower abundance of wild pollinators compared to honeybees in 2019 than in 2020 (54% of the visits vs, 31%, respectively), as wild insects are considered to be more effective than honeybees for crop pollination [9–11]. Contrary to the low fruit set, the seed set was in general relatively high, a pattern also found by other authors [54, 58], who suggested that carob trees selectively abort flowers with a smaller pollen load.

We expected an increase in fruit and seed production per female tree related to natural habitats [12, 15, 17, 18], and in orchards under ecological farming [21–24]. However, we did not find direct effects of these two variables on the number of fruits and seeds produced per female tree. We also expected a hump-shaped relationship between the proportion of male trees and fruit and seed production related to density-dependent competitive processes between sexes [34, 64]. As expected, local male-to-female ratio influenced the number of fruits and seeds produced per female tree following a hump-shaped relationship, being the yield per female maximized when the orchards had around 0.6-0.7 males/female tree, whereas the total production at the orchard level was maximized with 20-30% of males from the total. We must note, though, that this hump-shaped relationship was mainly driven by a study orchard with a male-to-female ratio higher than 1.5, and that the quadratic shape of this relationship was lost if that study orchard was not included in the analyses. Although sex ratio of wild carob trees in natural communities is approximately 1:1 [42, 90], which is the common sex ratio of diecious trees [42, 91], males were very scarce in most of our study orchards. This unbalanced sex ratio in crop fields is due to extended grafting practices that favor females for production, and prevented us finding more orchards with ratios higher than 1 to corroborate the hump-shaped relationship between male-to-female ratio and carob production. In any case, this hump-shaped relationship can be easily explained by the classic density-dependent model proposed by [34], so that males might facilitate females when at low densities whereas compete with them when at high densities.

However, from the production point of view, the weight of seeds might be as least as important than the number of fruits and seeds produced. Indeed, as many of the most valuable products derived from carob come from the seed [45, 46], the production of heavy seeds is very important for producers to increase their profits [53]. We expected that seed weight would be positively related to natural habitat in the landscape and local practices that enhance pollinator diversity and abundance [4, 10, 12, 13]. Here, we showed that the weight of carob seeds increased with overall pollinator abundance, but also with compositions of pollinator communities related to conserved natural habitats (higher abundance of wild bees and butterflies). On the contrary, pollinator communities associated with more disturbed landscapes (high abundance of honeybees) negatively influenced seed weight. The benefits of surrounding natural habitat for agricultural production have been described in several crops (e.g., [92–94]), usually through increased visit of wild pollinators that enhance crop yields. Here we found that conserving natural landscapes has clear benefits on carob seed weight by shaping more effective wild pollinator communities. These results agree with other studies also showing a positive effect of wild pollinator visits on seed weight (e.g., in soybean by [17], in soil rape by [95], or in sunflowers by [96]). Indeed, wild insects are described as more effective pollinators than honeybees for several crop species, as in honeydew melon in Israel [9], alfalfa in South Africa [97] or in several other species around the world as for instance almond, buckwheat, cotton, cherry, coffee, and kiwifruit (reviewed in [10]).

Seed weight was not only influenced by the pollinator community, but also by the male-to-female ratio of trees in the study orchards. While the number of carob seeds produced increased with the increasing male-to-female ratio, the weight of the seeds decreased. This opposite effect of the sex ratio on the quantity and quality of seeds might be explained by the well-known trade-off between seed number and size [98, 99], that arise from the limited amount of plant resources to invest in reproduction. The intra-specific trade-off between number and size (weight) has been described for multiple species (e.g., [100–103]). In crops, this trade-off is especially known in fruit trees, such as kiwifruit [104], apple [105], or peach [106], and is one of the reasons for pruning branches, as the reduction in the number of potential fruits increases the size of the remaining ones. In carob trees, the production of heavier seeds is of interest to farmers to increase their economic profits, as nowadays seeds are the most profitable part of the carob fruits (5-7€/kg in 2020, 7-20€/kg in 2021; 17-27€/kg in 2022) [53]. Here, we estimated that orchard-level maximum total yield in terms of weight occurs with 20-30% of males in the field. Therefore, implementing strategies for natural habitat and wild pollinator conservation and maintaining adequate male-to-female ratios will translate in a higher market value and increasing farmer’s profits [14, 95].

## Conclusions

Here we showed that natural landscapes promote wild pollinator communities (mostly wild bees) in carob tree orchards, while disturbed landscapes held primarily managed honeybees. Ecological farming tended to enhance pollinator abundance, while the carob tree male-to-female ratio also enhanced the abundance of pollinators at the orchard level, and modulated the production of fruit and seeds. Wild pollinator communities associated to conserved landscapes increased the weight of seeds, which are the most profitable part of carob production. For that, an adequate design of orchards with 20-30% of males, closely surrounded by natural habitats and subjected to ecological practices is important to optimize yields and profits.

## Acknowledgements

We are very grateful to the owners of the study orchards who allowed us working on their properties. This study was supported by the project CGL2017-89254-R, financed by the Spanish Ministry of Science and Innovation, FEDER Funds and the Spanish State Research Agency, and by the project PRPPID2020-117863RB-I00, financed by the Spanish Ministry of Science and Innovation and the Spanish State Research Agency (MCIN/AEI/10.13039/501100011033). CGM was supported by a FPI predoctoral contract financed by the Spanish Ministry of Economy and Competitiveness, the Spanish Research Agency, and European Social Funds (FPI PRE2018-083185, Call 2018). IMEDEA is an accredited “Maria de Maeztu Excellence Unit” (Grant CEX2021-001198, funded by MCIN/AEI/10.13039/501100011033) (period 2023-2027).

## Data availability

Data will be archived in DRYAD upon article acceptance.

## Authors’ contributions

**Carmelo Gómez-Martínez:** Methodology, Formal Analysis, Investigation, Resources, Data Curation, Writing – Original Draft, Writing – Reviewing & Editing, Visualization; **Miguel Ángel González-Estévez:** Investigation, Resources, Data Curation, Writing – Reviewing & Editing; **Indradatta deCastro-Arrazola:** Investigation, Writing – Reviewing & Editing; **Peter Unglaub:** Data Curation, Writing – Reviewing & Editing; **Amparo Lázaro:** Conceptualization, Methodology, Formal Analysis, Investigation, Writing - Original Draft; Writing – Reviewing & Editing.

